# Assessing the relative contributions of different *Arabidopsis thaliana* stigma factors to pollen hydration

**DOI:** 10.64898/2026.07.15.738742

**Authors:** Paula K.S. Chadic, Raymond K.M. Liu, Daphne R. Goring

**Author notes:** Corresponding Author: Daphne Goring. Equal contributions and first authorship (Paula K.S. Chadic, Raymond K.M. Liu).

## Abstract

Following pollination, *Arabidopsis* pollen grains rapidly hydrate through the transfer of water from the stigma to the pollen. Several stigma regulators of pollen hydration have been identified, and the corresponding mutants generally support milder defects in wildtype Col-0 pollen hydration, signifying the involvement of other unidentified factors in this process. Here, we uncovered a role for the stigma-specific mechanosensitive channel gene, *MscS-Like 7* (*MSL7*), in supporting pollen hydration. While the *msl7* mutant stigmas were found to support reduced hydration of wildtype Col-0 pollen, the phenotype was quite mild and very similar to that observed for other published pollen hydration mutants. Thus, we conducted a detailed comparison of different pollen hydration mutants on the stigma side (receptor kinases, PIP aquaporins) and pollen (PCP-Bs, MLS8) to compare pollen hydration mutant phenotypes and look for any additive effects of combining different mutants. Overall, all combinations resulted in the same mild hydration defect with no additional reductions in pollen hydration and no impact on pollen germination. This is in contrast to that observed for self-incompatible (SI) pollen from a transgenic *Arabidopsis* SI Col-0 line which shows very little pollen hydration and no pollen germination as part of the SI pollen rejection response. Together, these findings suggest that the regulation of compatible pollen hydration is quite complex and that there are likely other unknown mechanisms involved.

**Key Message:** The same mild pollen hydration defect is observed across very different *Arabidopsis* stigma mutants and it does not prevent pollen germination and pollen tube growth.

## Introduction

In *Arabidopsis*, reproductive success is determined by interactions between the male pollen and the female pistil. These interactions begin immediately after pollination, where pollen grains released from the anthers are deposited onto the unicellular hair-like projections on the surface of the stigma, known as the stigmatic papillae. Following pollen adhesion to a stigmatic papilla, compatible pollen signals are recognized and this initiate cellular events in the stigmatic papilla to facilitate the delivery of water to the desiccated pollen (reviewed in (Abhinandan et al. 2022; Bordeleau et al. 2022; Dickinson 1995; He et al. 2025). Rapid changes to the vacuolar morphology have been observed in the stigmatic papilla following pollination and thought to be linked to water transfer from the stigmatic papilla to the pollen (Fukushima et al. 2025). The stigmatic papilla supports pollen hydration to enable sufficient water uptake by the pollen to proceed to the next stage of pollen germination (Dickinson 1995; Hiroi et al. 2013; Rozier et al. 2020). Following germination, the pollen tube grows through the stigmatic papilla cell wall to the base where it enters the pistil reproductive tract and grows toward an ovule for fertilization (reviewed in (He et al. 2025; Zhong et al. 2025).

The pollen coat on the surface of the Arabidopsis pollen grain harbors small pollen coat proteins (PCPs) that can act as signaling peptides for pollen recognition (Wang et al. 2023). Three *PCP-Bs* (*α, β* and *γ*) were identified to promote pollen hydration as single and combination *pcp-b* mutant pollen displayed slower pollen hydration rates (Wang et al. 2017). These *PCP-Bs* are perceived by two sets of receptor kinases in the stigma: CrRLK1L members, FERONIA (FER) and ANJEA (ANJ), and S-DOMAIN RECEPTOR LIKE KINASE 28 (SD-RLK28). FER and ANJ function to both prevent and promote pollen hydration (Liu et al. 2021; Liu et al. 2025). In the absence of pollen, the FER-ANJ complex perceives stigma RALF22/33 peptides and activates the production of reactive oxygen species (ROS) in the stigmatic papilla to inhibit the hydration of unwanted pollen. In support of this, wildtype displays faster pollen hydration rates on the corresponding mutant stigmas (Liu et al. 2021). Following pollen contact, *PCP-Bγ* was proposed to outcompete RALF22/33 to bind to the FER-ANJ complex and cause a reduction in the stigmatic ROS levels to enable pollen hydration (Liu et al. 2021). More recently, three type 2C phosphatases were implicated as negative regulators of FER where their actions also reduced stigmatic ROS levels to enhance pollen hydration (Cheng et al. 2025). As well, FER-ANJ were implicated in the regulation of plasma membrane H^+^-ATPase (AHA) transporters which in turn were connected to the regulation of PIP aquaporins in the stigma (Liu et al. 2025). RALF22/33 binding to FER-ANJ in the stigma was proposed to cause AHA phosphorylation on an inhibitory site resulting in a lower cytosolic pH that leads to a closure of the PIP channels. Post-pollination, *PCP-Bγ* binding to FER-ANJ was proposed to result in AHA phosphorylation on activation sites, leading to a rise in the cytosolic pH and opening of the PIP channels to transport water out to the pollen for hydration (Liu et al. 2025). Aquaporins in the stigma have also been previously implicated in pollen hydration by Windari et al. (2021).

PCP-Bβ was also shown to bind to SD-RLK28, and *sd-rlk28* mutant stigmas supported slower pollen hydration and delayed pollen tube growth, though stigmatic ROS changes were unaffected (Guo et al. 2023). Lastly, a third set of receptor kinases (RKs), members of the Leucine-Rich Repeat Malectin (LRR-MAL) RKs, the linked *RECEPTOR-LIKE KINASE IN FLOWERS 1* (RKF1), and its paralogs, *RECEPTOR-LIKE KINASE IN FLOWERS 1* (*RKFL1*), *RKFL2*, and *RKFL3*, were shown to be required in the stigma to promote pollen hydration. The CRISPR/Cas9 mediated deletion of this gene cluster (*rkfΔ* mutants) resulted in slower pollen hydration rates on the mutant stigmas (Lee et al. 2024; Lee and Goring 2021). Collectively, the identification of FER-ANJ, SD-RLK28, and the RKF1/RKFLs highlights the involvement of multiple receptor kinases in the stigma to regulate pollen hydration. Secretion in the stigmatic papillae is also connected to pollen hydration with the exocyst and SNARE complexes being required for this process (Macgregor et al. 2023; Safavian and Goring 2013; Safavian et al. 2015; Samuel et al. 2009). This secretion may be responsible for transporting proteins to the stigmatic papilla plasma membrane to promote pollen hydration.

Here we have identified the *MscS-like 7* (*MSL7)* gene as another stigma regulator of pollen hydration. *MSL7* was investigated because its closest paralog, *MSL8*, was previously identified to be a pollen-specific gene required for osmoregulation during pollen hydration *in vitro* (Hamilton et al. 2015). *MSL7* is tandemly-linked to *MSL8* and has a very restricted pattern of expression with high levels in the stigma and lower levels in pollen tubes following *in vitro* growth through the stigma and style (Wang et al. 2022). During our investigations on *MSL7*, we noticed that various stigma mutants generally supported the same mild hydration defect for wildtype Col-0 pollen. Thus, we used a standard assay to conducted a detailed analysis of different pollen hydration mutants on the stigma side (receptor kinases, PIP aquaporins) and pollen side (PCP-Bs, MLS8) to assess the degree of reduced pollen hydration across these mutants and to better understand their relative contributions to this process.

## Materials and Methods

### Plasmid construction and plant transformation

The CRISPR/Cas9 induced *msl7* mutations were generated as previously described (Doucet et al. 2019a; Doucet et al. 2019b; Wang et al. 2015). Target sites for deletion mutations in *MSL7* (At2g17000) were selected through CHOPCHOP (Labun et al. 2019), and the final pBEE401E vector contained two gRNAs located towards each end of the *MSL7* gene (Suppl. Table S2). Wildtype *A. thaliana* Col-0 plants were transformed through *Agrobacterium*-mediated gene transfer via floral dip, and plants were dipped twice with a 7-day interval to maximize transformation efficiency (Clough and Bent 1998). Seeds collected from these plants were stratified and germinated on soil. In the T1 generation, transformants were selected by BASTA screening, where 7-day old seedlings were sprayed with the BASTA herbicide for a total of three times with 2-day intervals. Transformants were also confirmed through the PCR detection of the BASTA resistance gene, and deletion mutations in *MSL7* were identified through PCR genotyping. These plants were then left to set seed, and the subsequent T2 generation plants were screened for homozygous *msl7* mutants resulting in two mutants, *msl7-2* and *msl7-3* (Suppl. Fig. S1).

### Plant materials and growth conditions

Seeds stocks used in this study were *rkfΔ-1,-2* (Lee and Goring 2021), *sd-rlk28* (SALK_065560 from ABRC; (Guo et al. 2023), *msl8-5* and the *msl7-1 msl8-6* (Wang et al. 2022), pcp-bɑ/β/γ seeds (Wang et al. 2017), and the different *pip* mutant combinations (Ceciliato et al. 2019; Schley et al. 2025). Higher order mutants were generated by crossing the *msl7-2* and msl7-3 mutants with *rkfΔ-1* and *rkfΔ-2* mutants, respectively. Plants that were heterozygous for the *msl7* and *rkfΔ* mutations in the F1 generation were identified and left to set seed, and plants that were homozygous for the *msl7* and *rkfΔ* mutations were identified in the T2. Finally, the F3 generation was screened to identify plants that were homozygous for both the mls7 and rkfΔ mutations. All mutants and genotyping primers used in this study are listed in Suppl. Table S3. The transgenic *Arabidopsis* SI Col-0 line was previously described (Macgregor et al. 2022; Zhang et al. 2019). All seeds were stratified in the dark at 4° C for at least 5 days before placement on Sunshine #1 soil with Plant Prod All Purpose 20-20-20 fertilizer for germination and growth. Plants were grown in chambers at 22° C with 16-h light/8-h dark, and with the humidity maintained below 50%.

### Pollen hydration and aniline blue staining

Post-pollination assays were conducted as described in Lee et al. (2020). In preparation for the assays, wildtype and mutant stage 12 flower buds were emasculated, wrapped in plastic wrap, then returned in the growth chambers. For the pollen hydration assay, at 24-h post-emasculation, pistils were collected and mounted on ½ MS plates. A single anther from an open flower was used to lightly pollinate the mounted pistils and images were captured at 0- and 10-min post-pollination using a Nikon sMz800 microscope. 10 pollen grains were randomly measured per pistil and this assay was repeated on 3 pistils per line, measuring a total of 30 pollen grains per group (*n* = 30). For the aniline blue staining assay, at 24-h post-emasculation, pistils were pollinated with a single anther from open flowers. At 4-h post-pollination, pistils were collected and stained as described in Lee et al. (2020). Aniline blue-stained pistils were mounted on a microscope slide with sterile water and flattened with a coverslip, and images were captured on 10X magnification using the brightfield and UV fluorescence settings on a Zeiss Axioskop2 Plus fluorescent microscope. For each pistil, 2-3 images were taken progressively moving down the transmitting tract to track the pollen tubes and the pollen tube front. A high resolution composite image for each pistil was then assembled in Adobe Photoshop using the pixel-by-pixel move tool to precisely align the images together. The pollen tube front was then measured by determining the distance between the base of the style and the pollen tube front using the built-in ruler tool. Measurements were taken from a horizontal line at the base of stigma to the pollen tube front (leading edge of the majority of growing pollen tubes in the transmitting tract; (Chae et al. 2009; Lee and Goring 2021). The pixels were converted to microns using a stage micrometer image taken at the same 10X magnification. For statistical analyses, SPSS (IBM) was used to perform one-way analysis of variance (ANOVA) with Tukey’s honest significant difference (HSD) post-hoc test. Cut-off value was specified as P < 0.05.

## Results

### A role for *MSL7* in the stigma to support wildtype Col-0 pollen hydration

Previously, *MSL7* was identified in searches for genes displaying stigma-enriched expression (Doucet et al. 2019a; Doucet et al. 2019b), and found to be highly expressed in stigma and stigmatic papillae RNA-seq datasets (Suppl. Table S1 (Gao et al. 2018; Klepikova et al. 2016; Wang et al. 2022). To investigate potential functions for *MSL7* during pollen-stigma interactions, CRISPR/Cas9 was used to generate two *Arabidopsis MSL7* deletion mutants resulting in *msl7-2* and *msl7-3* (Suppl. Fig. S1). Stigmas from these *mls7* mutants were then used in pollen hydration assays to assess whether *MSL7* is required in the stigma to support wildtype pollen hydration (Fig. 1). The *msl7-1 msl8-6* mutant (Wang et al. 2022) was also included to test for any potential contributions of the tandemly-linked, pollen expressed *MSL8* gene (Suppl. Fig. S1). Wildtype Col-0 pollen grains were applied to the mutant stigmas, and pollen grain diameters were measured, as a proxy for hydration, at 0- and 10-min post-pollination (Lee et al. 2020). Stigmas from the *msl7-2, -3* mutants supported significantly reduced levels of Col-0 pollen hydration at 10-min post pollination when compared to the wildtype Col-0 stigmas (Fig. 1a). The same level of reduced Col-0 pollen hydration was also observed in the *msl7-1 msl8-6* double mutant indicating that the *msl7* mutant stigma phenotype is independent of *MSL8.* As such, these findings indicate that *MSL7* functions in the stigma to support wildtype pollen hydration.

**Fig. 1.**
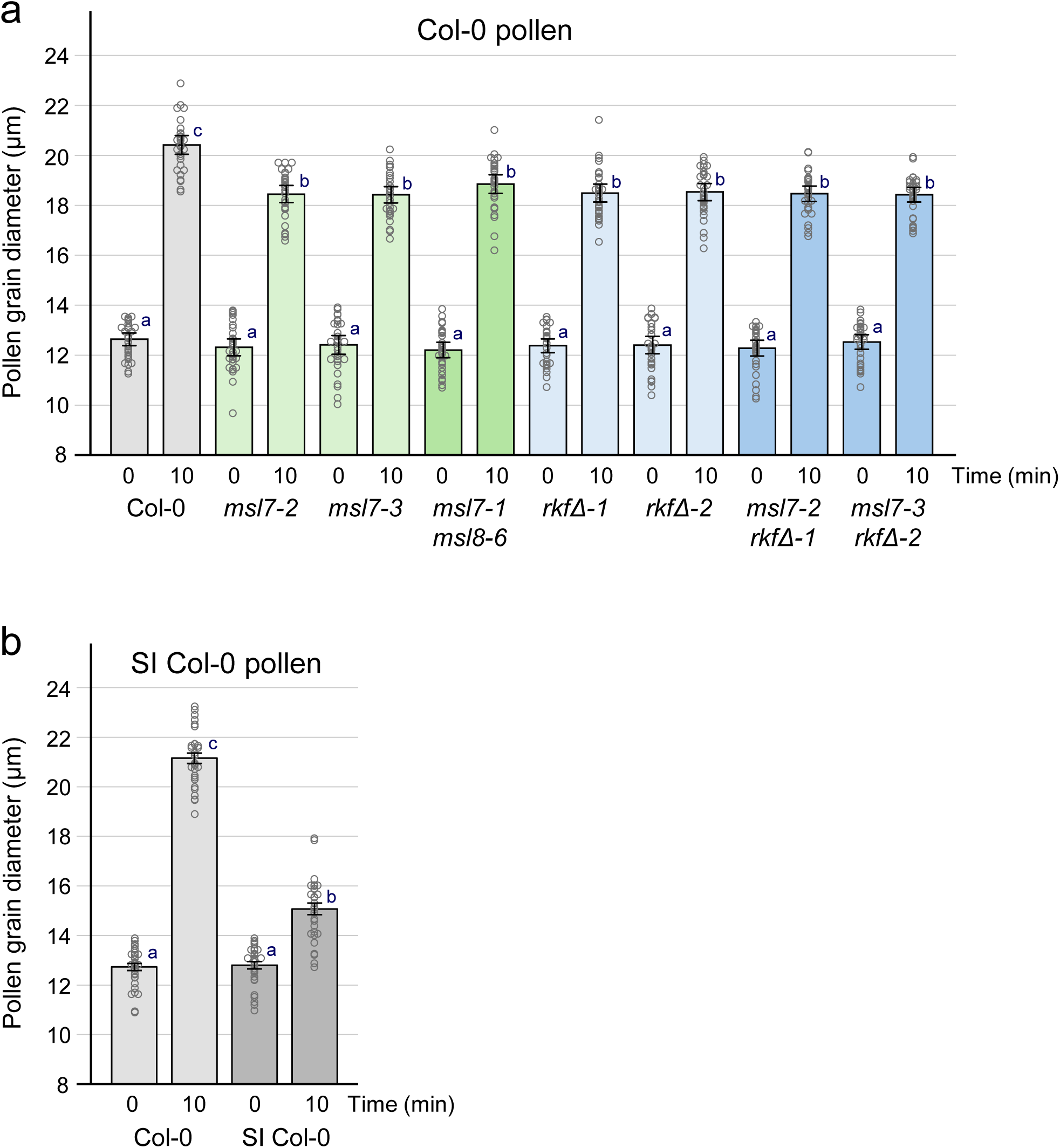
Col-0 Pollen hydration assay on various *msl* and *rkfΔ* mutant stigmas. **a** The hydration of Col-0 pollen grains was measured on compatible stigmas from Col-0, *msl7*, *msl7 msl8*, *rkfΔ,* and *msl7 rkfΔ* mutants with Col-0 pollen hydration on Col-0 stigmas as a positive control. Reduced pollen hydration was observed on the mutant stigmas. **b** SI Col-0 pollen hydration on SI Col-0 stigmas is shown as a comparison of how strongly pollen hydration can be inhibited. Pollen diameters were measured at 0- and 10-min post-pollination. Data are shown as bar graphs ± SE with all the data points displayed. *n* = 30 pollen grains per line. Letters represent statistically significant groupings of *P* <0.05 based on a one-way ANOVA with a Tukey-HSD post-hoc test.

The decreased hydration of Col-0 pollen observed with the *msl7* mutant stigmas at 10 min post-pollination was also equivalent to that observed in the previously characterized *rkfΔ-1, -2* mutants which carry a single deletion of the four tandemly-linked *RKF1/RKFLs* genes (Fig. 1a; (Lee and Goring 2021). These mutants were combined to determine if *MSL7* and the *RKF1/RKFLs* functioned together in the stigma to promote pollen hydration. However, stigmas from both the *msl7-2 rkfΔ-1* and the *msl7-3 rkfΔ-2* mutants did not support an additive decrease in wildtype Col-0 hydration at 10-min post-pollination (Fig. 1a). As a result, the average Col-0 pollen diameters at 10 min post-pollination were not significantly different on stigmas from the single *msl7-2, -3* mutants, the *msl7-1 msl8-6* double mutant and the *msl7 rkfΔ* mutants (Fig. 1a). Despite the significant decreases in hydration observed for Col-0 pollen on these mutant stigmas, this was still a relatively mild phenotype. At 10-min post-pollination, Col-0 pollen had an average diameter of ∼18.5 µm on the mutant stigmas compared to ∼20.4 µm on Col-0 stigmas (Fig. 1a), and this did not prevent pollen germination. As a comparison, pollen hydration is strongly inhibited by self-incompatibility (Fig. 1b), and this effectively prevents pollen germination (Abhinandan et al. 2022; Goring et al. 2023; Rozier et al. 2020). At 10 min post-pollination, SI pollen from the transgenic *Arabidopsis* SI Col-0 line had an average diameter of ∼15 µm on the SI-Col-0 stigma compared to ∼21 µm on Col-0 stigmas (Fig. 1b; (Macgregor et al. 2022; Zhang et al. 2019).

### Impact of combining pollen and stigma mutants on pollen hydration

Since the *msl7 rkfΔ* combined mutants did not display any additive effects on reducing pollen hydration, we were interested to see if pollen mutants with hydration defects would display further decreases in hydration when placed on these mutant stigmas. As well, we carried out a direct comparison to other published stigma mutants to determine their relative effects on pollen hydration when tested alongside the *msl7 rkfΔ* mutant stigmas (Table 1). Previously, the triple *pcp-bα/β/γ* mutant pollen were shown to have slower hydration rates on Col-0 stigmas compared to wildtype Col-0 pollen (Wang et al. 2017), and pollen PCP-Bs were implicated in regulating the FER-ANJ and SD-RLK28 receptor kinases in the stigma to promote pollen hydration (Guo et al. 2023; Liu et al. 2021; Liu et al. 2025). Thus, hydration of these triple *pcp-bα/β/γ* mutant pollen grains was examined when placed on the stigma mutants compared to wildtype Col-0 stigmas. With MSL8 having a role in regulating pollen hydration under hypoosmotic conditions *in vitro* (Hamilton et al. 2015), *msl8-5* mutant pollen hydration was also included. Pollen hydration of these two different pollen mutants were conducted on stigmas from wildtype Col-0 and the *msl7-2/3*, *rkfΔ-1/2, msl7-2/3 rkfΔ-1/2,* and *msl7-1 msl8-6* mutants (Fig. 2). Col-0 pollen hydration on Col-0 stigmas was used as the control for wildtype pollen hydration rates. The *pcp-bα*/*β*/*γ* mutant pollen on the wildtype Col-0 stigmas exhibited a mild reduction in pollen hydration at 10 min post-pollination as expected (Fig. 2a; (Wang et al. 2017). Again, there were no additional decreases in pollen hydration at 10 min post-pollination for the *pcp-bα*/*β*/*γ* mutant pollen placed on stigmas from the *msl7-2/3*, *rkfΔ-1/2,* and *msl7-2/3 rkfΔ-1/2* mutants, with the *pcp-bα*/*β*/*γ* mutant pollen essentially displaying the same level of hydration as that observed on the Col-0 stigmas (Fig. 2a). The *msl8-5* mutant pollen did not have any reduced pollen hydration at 10 min post-pollination on Col-0 stigmas as previously shown (Hamilton et al. 2015), and displayed the same phenotype of a mild reduction in pollen hydration as Col-0 pollen and *pcp-bα/β/γ* mutant pollen on the various stigma mutants (Fig. 2b). Lastly, stigmas from the *sd-rlk28* mutant which was previously reported to support reduced pollen hydration levels (Guo et al. 2023) were tested alongside the *msl7-2 rkfΔ-1* stigmas in pollen hydration assays with pollen from Col-0, the *pcp-bα/β/γ* mutant, and the *msl8-5* mutant. The wildtype and mutant pollen grains showed similar mild reductions in hydration on the *sd-rlk28* and *msl7-2 rkfΔ-1* mutant stigmas. The decreases in pollen hydration at 10 minutes post-pollination for these mutants were significantly different from Col-0 pollen hydration on Col-0 stigmas, but not significantly different from each other (Fig. 3). Thus, the two different pollen mutants did not display any further decreases in pollen hydration at 10 min post-pollination on the *msl7-2 rkfΔ-1* and *sd-rlk28* mutant stigmas (Fig. 1 and 2).

**Fig. 2.**
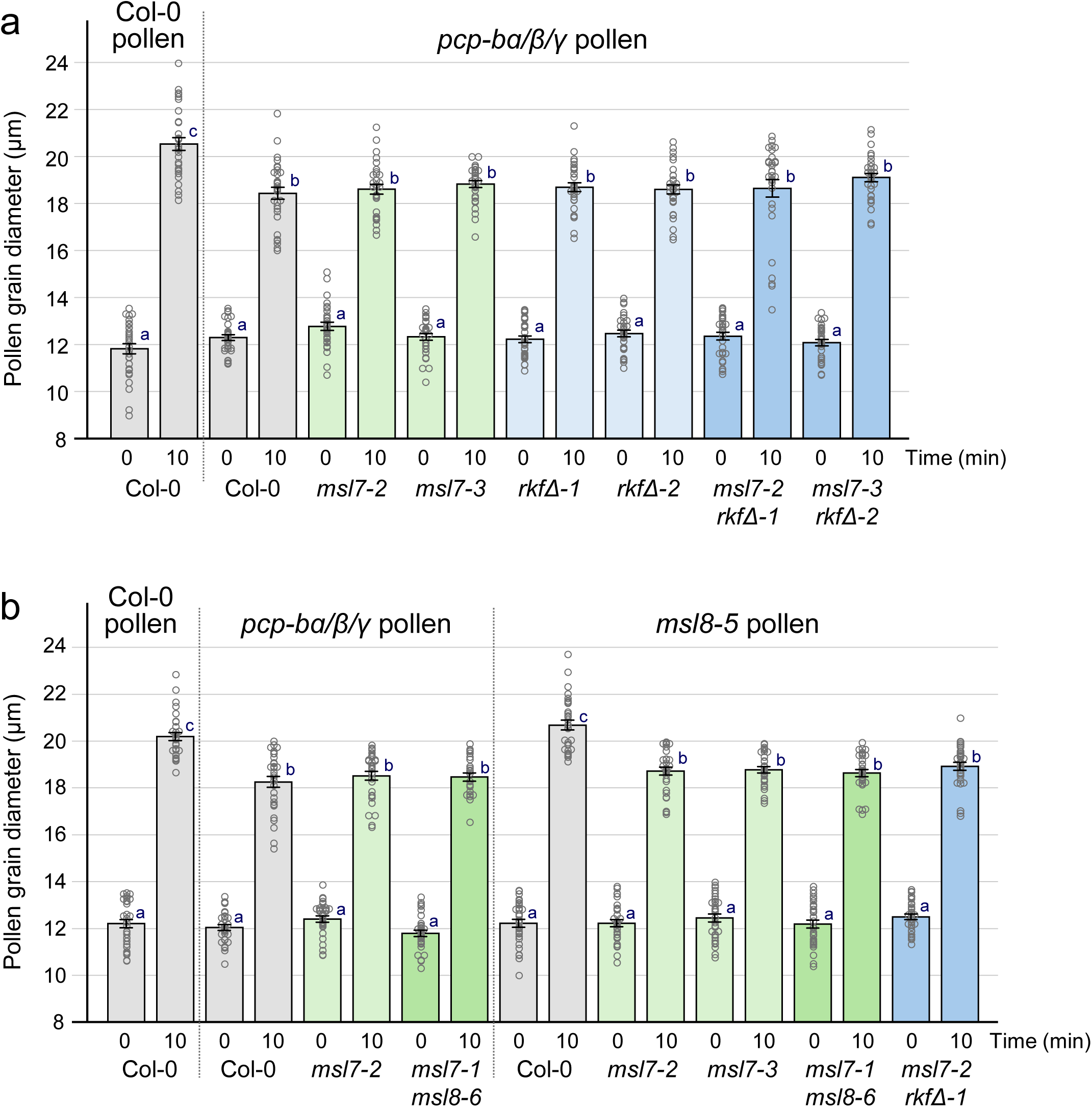
Comparing *pcp-bɑ/β/γ* and *msl-8-5* pollen hydration on various *msl* and *rkfΔ* mutant stigmas. The hydration of *pcp-bɑ/β/γ* and *msl-8-5* pollen grains was measured on compatible stigmas from Col-0, *msl7*, *msl7 msl8*, and *msl7 rkfΔ* mutants with Col-0 pollen hydration on Col-0 stigma as a positive control. **a** Similar levels of reduced *pcp-bɑ/β/γ* pollen hydration were observed on Col-0 and the mutant stigmas. **b** While *pcp-bɑ/β/γ* pollen hydration was reduced on Col-0 stigmas, the *msl-8-5* pollen showed wildtype levels of pollen on Col-0 stigmas. In contrast, their levels of hydration on the different stigma mutants were not significantly different. Pollen diameters were measured at 0- and 10-min post-pollination. Data are shown as bar graphs ± SE with all the data points displayed. *n* = 30 pollen grains per line. Letters represent statistically significant groupings of *P* <0.05 based on a one-way ANOVA with a Tukey-HSD post-hoc test.

**Fig. 3.**
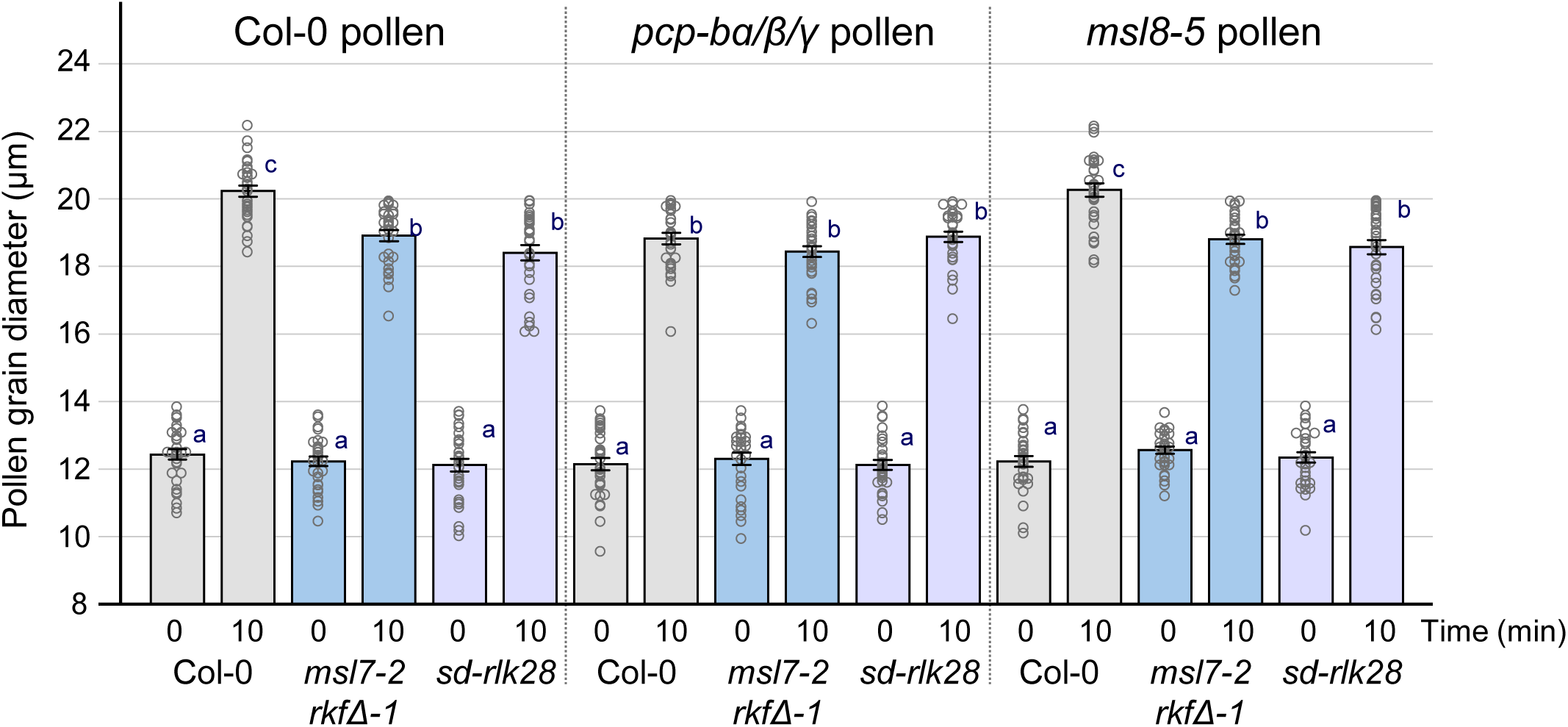
Comparing *pcp-bɑ/β/γ* and *msl-8-5* pollen hydration on the *sd-rlk28* mutant. The hydration of *pcp-bɑ/β/γ* and *msl-8-5* pollen was measured on compatible stigmas from Col-0, *msl7-2 rkfΔ-1*, and *sdrlk28-1* mutants. The *sdrlk28-1* stigmas supported the same level of reduced pollen hydration for pollen from Col-0, *pcp-bɑ/β/γ* and *msl-8-5 plants* as the *msl7-2 rkfΔ-1* mutant stigmas. Pollen diameters were measured at 0- and 10-min post-pollination. Data are shown as bar graphs ± SE with all the data points displayed. *n* = 30 pollen grains per line. Letters represent statistically significant groupings of *P* <0.05 based on a one-way ANOVA with a Tukey-HSD post-hoc test.

**Table 1.**
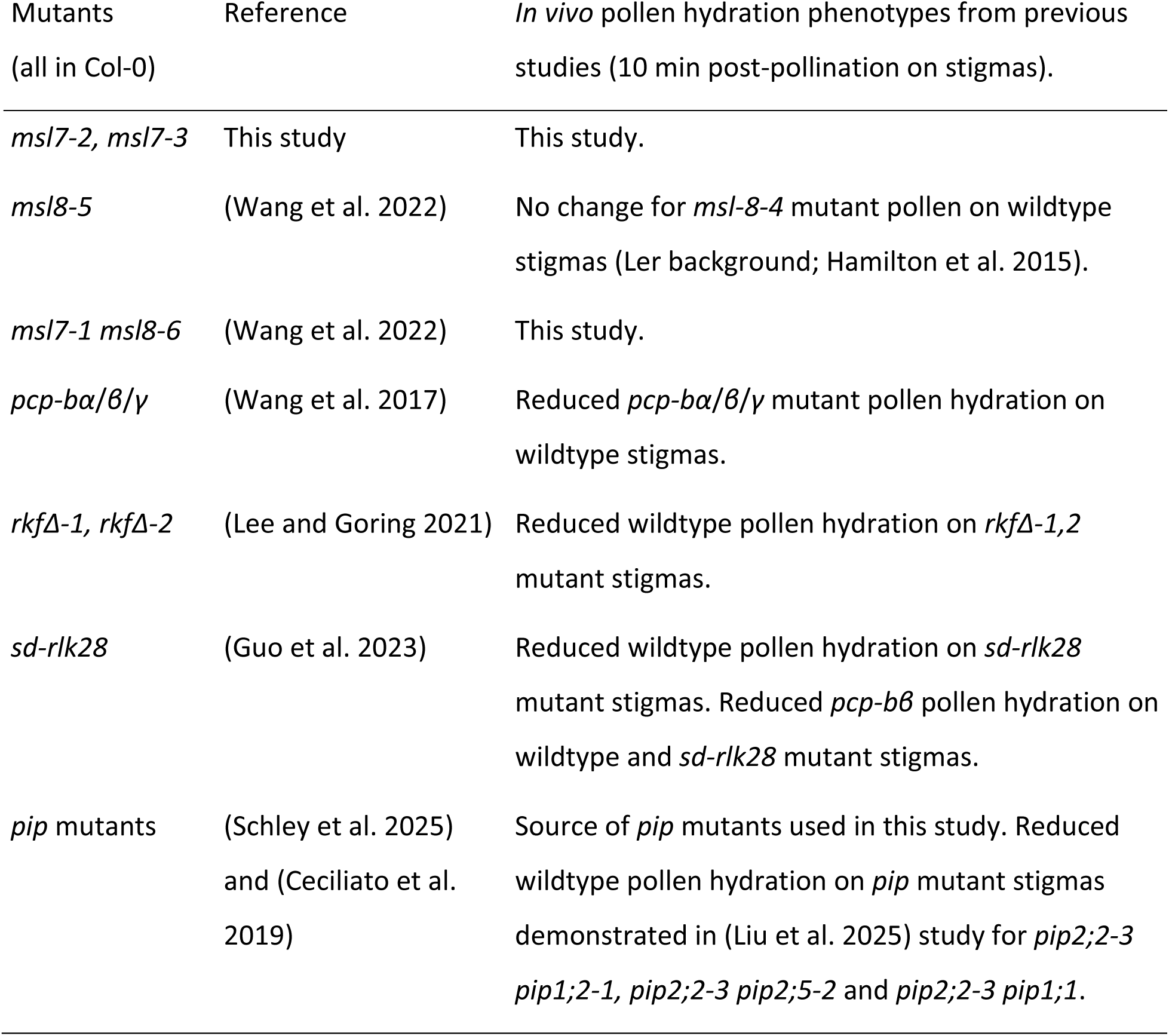
Mutants used in this study.

Plasma membrane intrinsic proteins (PIPs) have been implicated as the aquaporin water channels in the stigmatic papillae to promote water export to pollen (Liu et al. 2025; Windari et al. 2021), and so *pip* stigma mutants were directly compared to mutant stigmas from *sd-rlk28* mutant and the *rkfΔ-1 nilr2rir1Δ-1 lik1-5* mutant (which supports the same Col-0 hydration phenotype as *rkfΔ-1*; (Lee et al. 2024) to determine their relative impact on Col-0 pollen hydration (Fig. 4). PIP1;2 and the ER aquaporin, SIP1;1, were first reported as factors in this process (Windari et al. 2021). More recently, PIP1;2, PIP2;2, PIP1;1 and PIP2;5 were identified as stigma factors to promote pollen hydration (Liu et al. 2025). These four PIPs were tested as double mutant combinations of *pip2;2* with *pip1;2*, *pip1;1* or *pip2;5* and all three double mutants supported the same reduced level of pollen hydration. Lui et al. (2025) also generated RNAi lines targeting all 13 PIPs using a stigma-specific promoter, and stigmas from these RNAi lines did not show any further decrease in Col-0 pollen hydration at 10 min post-pollination compared to that observed for the double *pip* stigma mutants. Here we tested a quadruple *pip* mutant, *pip1;1 pip1;2 pip2;1 pip2;2* (Ceciliato et al. 2019; Schley et al. 2025), that overlaps with the previously tested *pip* mutant stigmas (Liu et al. 2025) in pollen hydration assays. An analysis of transcriptome datasets showed that among the thirteen Arabidopsis *PIPs*, *PIP1;2* and *PIP2;2* show the highest expression in a stigmatic papilla transcriptome dataset, and *PIP1;2* also shows the highest expression in a stigma transcriptome dataset (Suppl. Table S1; (Gao et al. 2018; Klepikova et al. 2016). Wildtype Col-0 pollen hydration was measured at 10 min post-pollination on stigmas from the quadruple *pip* mutant alongside the single *pip1;2-2*, *pip2;1-2* and *pip2;2-3* mutants and the *sd-rlk28* and *rkfΔ-1 nilr2rir1Δ-1 lik1-5* mutants (Fig. 4). The single *pip1;2-2*, *pip2;1-2* and *pip2;2-3* mutant stigmas supported a slight but significant decrease in wildtype

**Fig. 4.**
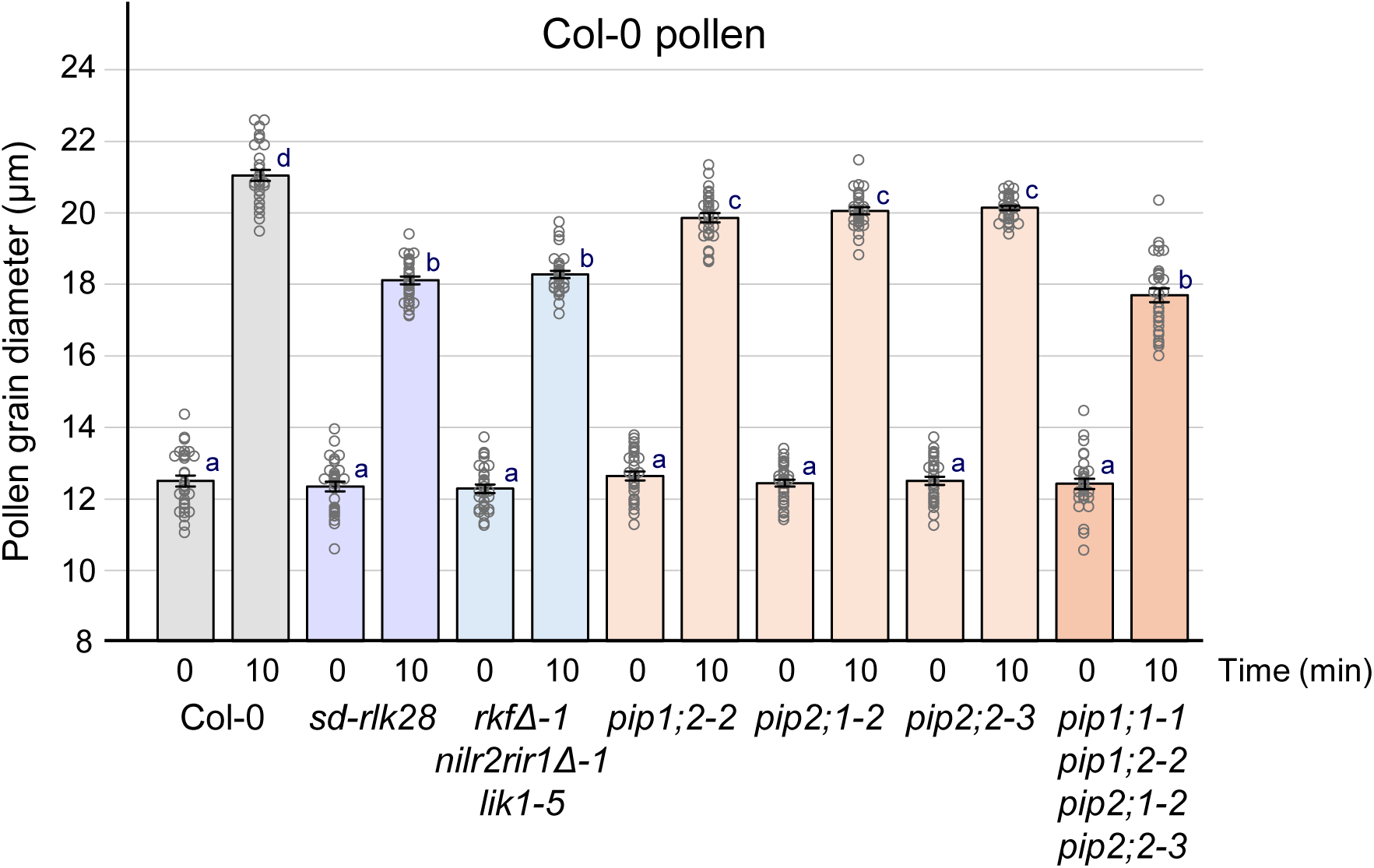
Col-0 pollen hydration on different *pip* mutants. The hydration of Col-0 pollen was measured on compatible stigmas from Col-0 and the various *pip* mutants. The quadruple *pip* mutant stigmas supported the same level of reduced Col-0 pollen hydration at 10 min post-pollination as that seen for the *sdrlk28-1* and the *rkfΔ-1 nilr2rir1Δ-1 lik1-5* stigmas. In comparison, Col-0 pollen hydration on the single *pip* mutant stigmas was higher but still reduced compared to that seen for Col-0 stigmas. Pollen diameters were measured at 0- and 10-min post-pollination. Data are shown as bar graphs ± SE with all the data points displayed. *n* = 30 pollen grains per line. Letters represent statistically significant groupings of *P* <0.05 based on a one-way ANOVA with a Tukey-HSD post-hoc test.

Col-0 pollen hydration compared to Col-0 pollen on Col-0 stigmas while the quadruple *pip1;1-1 pip1;2-2 pip2;1-2 pip2;2-3* mutant stigmas supported a further significant decrease in Col-0 pollen hydration (Fig. 4). At 10-min post-pollination, Col-0 pollen had an average diameter of ∼19.9-20.2 µm on stigmas from the single *pip* mutants and ∼17.7 µm on quadruple *pip1;1-1 pip1;2-2 pip2;1-2 pip2;2-3* mutant stigmas compared to ∼21.1 µm on wildtype Col-0 stigmas. The decrease in wildtype Col-0 pollen hydration on the quadruple *pip1;1-1 pip1;2-2 pip2;1-2 pip2;2-3* mutant stigmas was not significantly different to that observed for Col-0 pollen on the *sd-rlk28* and *rkfΔ-1 nilr2rir1Δ-1 lik1-5* mutant stigmas (Fig. 4). Thus, quadruple *pip* mutant stigmas supported the same reduced level of pollen hydration as observed for all the other mutant combinations tested in this study.

### Col-0 and *pcp-bα/β/γ* mutant pollen tube growth on different stigma mutants

While the degree of Col-0 and *pcp-bα/β/γ* mutant pollen hydration on the various mutant stigmas was significant reduced at 10 minutes post-pollination in comparison to Col-0 pollen hydration on Col-0 stigmas, it was not sufficient to block pollen germination. As a result, abundant pollen tubes were observed in pollinated pistils. However, *pcp-bα/β/γ* mutant pollen tubes were previously reported to have reduced growth lengths in wildtype Col-0 pistils at 4-hrs post-pollination (Guo et al. 2023; Wang et al. 2017) and so we followed up on this observation to assessed whether pollen tube growth lengths might be altered in the mutant pistils studied here (Fig 5). Wildtype Col-0 and *pcp-bα/β/γ* mutant pollen tube growth lengths were measured in pistils from wildtype Col-0 and three different mutants: *sd-rlk28*, *msl7-2 rkfΔ-1* and the *pip1;1-1 pip1;2-2 pip2;1-2 pip2;2-3* quadruple mutant (Fig. 5). The pollinated pistils were collected at 4 hrs post-pollination, and stained with aniline blue to visualize the pollen tubes and measure the pollen tube fronts (Chae et al. 2009; Lee and Goring 2021). The wildtype Col-0 pollen tube growth lengths did not vary in the different mutant pistils and were not significantly different from that observed in wildtype Col-0 pistils (Fig. 5a-d, i). The *pcp-bα/β/γ* mutant pollen tubes growth lengths were significantly shorter as previously reported (Guo et al. 2023; Wang et al. 2017), but there were no significance differences in the *pcp-bα/β/γ* mutant pollen tubes growth lengths across pistils from wildtype Col-0 and the *sd-rlk28*, *msl7-2 rkfΔ-1* and the *pip1;1-1 pip1;2-2 pip2;1-2 pip2;2-3* mutants (Fig. 5e-h, i). These results indicated that there were no additional effects on the growth of wildtype Col-0 and *pcp-bα/β/γ* mutant pollen tubes through the *sd-rlk28*, *msl7-2 rkfΔ-1* and the *pip1;1-1 pip1;2-2 pip2;1-2 pip2;2-3* mutant pistils.

**Fig. 5.**
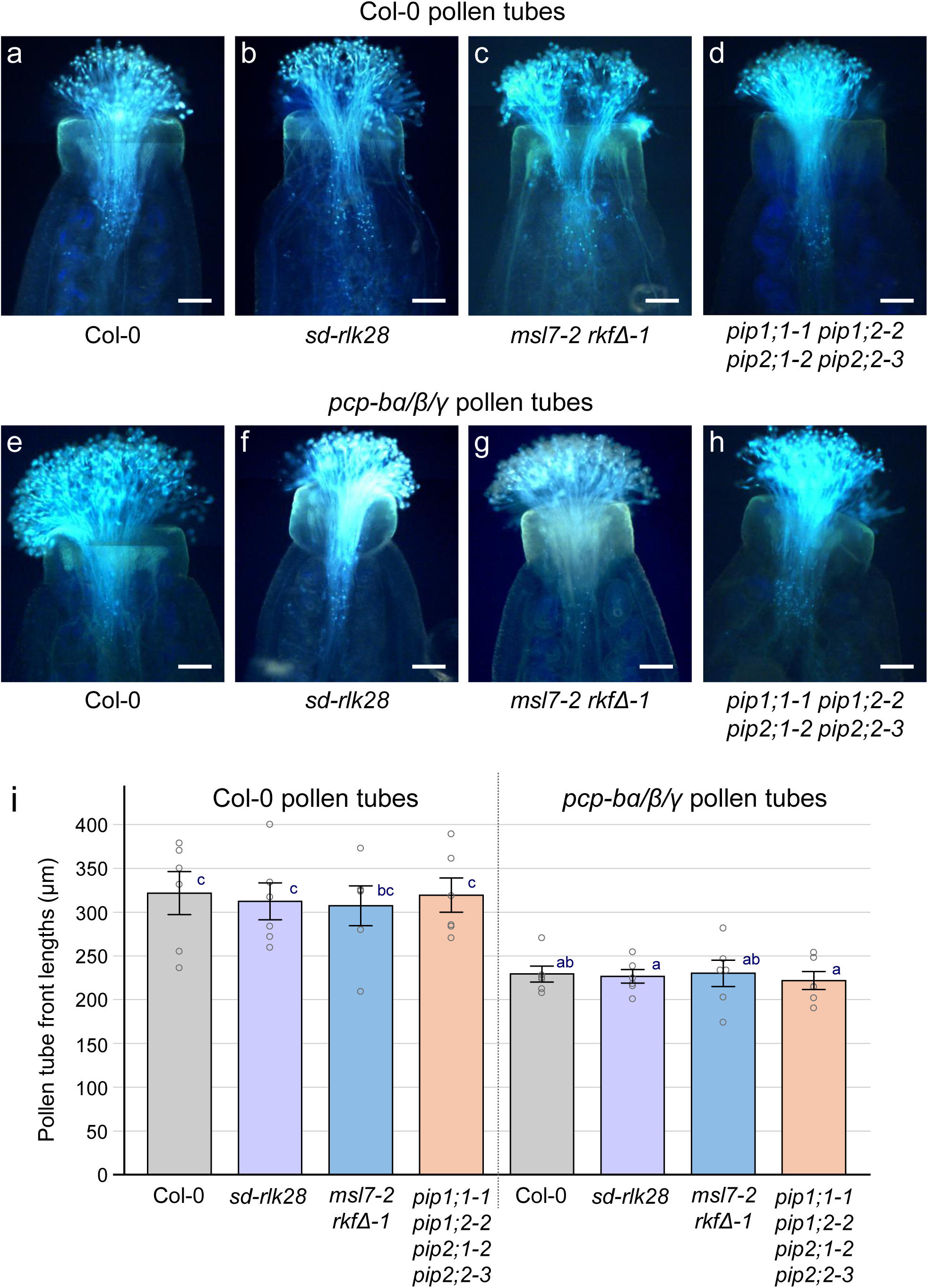
Growth of Col-0 and *pcp-bɑ/β/γ* pollen tubes on different mutant pistils at 4-h post-pollination. **a-h.** Representative aniline blue images of pollinated pistils at 4-h post pollination. Genotypes of the pistils and pollen tubes are indicated below and above, respectively. Shown are composite images assembled from overlapping images. Scale bars = 100 μm. **i.** Mean lengths of the Col-0 and *pcp-bɑ/β/γ* pollen tube fronts measured from the base of the styles. Data are shown as bar graphs ± SE with all the data points displayed. *n* = 6 pollinated pistils per line. Letters represent statistically significant groupings of *P* <0.05 based on a one-way ANOVA with a Tukey-HSD post-hoc test.

## Discussion

In this study, we investigated factors in the stigma responsible for promoting wildtype pollen hydration and identified *MSL7* as another stigma component involved in this process. We observed that the loss of *MSL7* in the stigma resulted in a mild but significant reduction in the degree of wildtype Col-0 pollen hydration at 10 min post-pollination (Fig. 1). Plasma membrane MSLs are proposed to be similar to the bacterial MscS channel and function as tension-gated channels to open in response to membrane tension and release osmolytes to reduce turgor pressure (Guichard et al. 2022; Hamilton et al. 2015; Naismith and Booth 2012). In this proposed role, the loss of stigma MLS7 may be more of an indirect effect on pollen hydration. Failure of the *msl7* mutant to reduce a hypothetical stigmatic papilla membrane tension following pollination could be linked to reduced aquaporin water transport caused by increased membrane tension (Caviglia et al. 2024; Ozu et al. 2023),

As the mild reduced pollen hydration phenotype on *msl7* mutant stigmas was similar to that observed for the *rkfΔ* mutant stigmas (Lee and Goring 2021), we investigated how widespread trait was by incorporating other published pollen hydration mutants to our analyses: the *sd-rlk28* stigma mutant (Guo et al. 2023), *pip* stigma mutants (Liu et al. 2025; Windari et al. 2021) and the *pcp-bα*/*β*/*γ* pollen mutant (Wang et al. 2017). This allowed for a direct comparison of the impact of these different mutants on pollen hydration, and it was through this comparison that we discovered that the mutants were all producing the same mild but significant reduction in pollen hydration at 10 min post-pollination. Furthermore, we found that combining different stigma mutants did not result in any additional decreases in pollen hydration rates at 10 minutes post-pollination. Finally, this reduced pollen hydration phenotype was consistent for mutant stigmas regardless of whether wildtype Col-0 or *pcp-bα*/*β*/*γ* mutant pollen grains were added (Fig. 1-4). Additionally, the reduced pollen hydration phenotypes at 10 minutes post-pollination did not affect the next steps of pollen germination and pollen tube growth (Fig. 5). Wildtype Col-0 pollen tubes grew the same distance at 4-hrs post-pollination in pistils from wildtype Col-0 and the stigma mutants indicating there was no impact of these mutants on pollen tube growth. The *pcp-bα*/*β*/*γ* mutant pollen tubes were previously reported to grow a shorter distance compared to wildtype pollen tubes (Guo et al. 2023; Wang et al. 2017) and this was also observed in this study (Fig. 5).

It is well known that the Brassicaceae stigma plays a key role in promoting hydration for compatible pollen and preventing the hydration of self-incompatible (SI) pollen (reviewed in (Abhinandan et al. 2022; Dickinson 1995; Zhang et al. 2026). As shown in Fig. 1, the SI pollen rejection response in the transgenic *Arabidopsis* SI Col-0 line results in a significant reduction on SI pollen hydration compared to compatible Col-0 pollen. So, with the stigma’s key role in regulating pollen hydration, one would predict that knocking out the regulators of compatible pollen hydration would recapitulate the degree of pollen hydration suppression observed with SI pollen. However, this is not the case and so the question arises as to why there is a gap between the degree of reduced pollen hydration for compatible pollen hydration mutants compared to the SI pollen hydration.

When compatible pollen lands on a stigma, it adheres to a stigmatic papilla and an interface (pollen ‘foot’) forms between the pollen and the stigmatic papilla, composed of lipids, proteins, sugars and phenolics derived from the pollen coat and the surface stigmatic pellicle. The pollen coat is strongly hydrophobic but changes take place in the pollen foot through the proposed action of enzymes, such as extracellular lipases (in the pollen coat and stigmatic pellicle), to form hydrophilic channels for water transport from the stigma to the pollen grain, and this water movement is driven by the osmotic differential between the desiccated pollen grain and the stigmatic papilla (Dickinson 1995; Lyu and Liang 2025; Updegraff et al. 2009; Wang et al. 2023). Detailed profiles on *Brassica* and *Arabidopsis* compatible pollen hydration show that there are two phases to hydration. First, there is a high hydration rate leading to an almost fully hydrated pollen grain at 10 min post-pollination for *Arabidopsis* pollen (Rozier et al. 2020) and at 30-40 min post-pollination for *Brassica* pollen (Dickinson 1995; Hiroi et al. 2013). The following second phase has a much lower hydration rate and *Arabidopsis* pollen germinates a pollen at ∼ 20 min post-pollination, though the timing can be quite variable (Rozier et al. 2020). For *Brassica* pollen, there can be some increase or loss in pollen diameter in the second phase, and pollen will germinate a pollen tube at ∼40-50 min post-pollination (Dickinson 1995; Hiroi et al. 2013). In contrast to the compatible pollen hydration profiles, *Brassica* and *Arabidopsis* SI pollen typically showed only a small amount of hydration in the first phase, with very little increase in in the second phase with no pollen tube germination (Dickinson 1995; Hiroi et al. 2013; Rozier et al. 2020).

Focusing on the stigma-side, there are several possible explanations that may contribute to the pollen hydration gap we observed between SI pollen hydration and the compatible pollen hydration mutants. First, there may be other components and/or homologues that are still functioning to promote compatible pollen hydration; for example, additional stigma expressed PIPs may need to be knocked out to further impact pollen hydration. What was striking in this study was that all the different combinations tested resulted in the same level of reduced pollen hydration, and so it poses the possibility that other processes may be regulating pollen hydration. Dickinson (1995) proposed that before the high hydration rate phase, there is a variable period of time where hydraulic continuity is set up in the pollen foot between the pollen and stigmatic papilla and some water is transferred by capillary action. Perhaps, there are passive water transfer processes involved in the early phase of pollen hydration, such as water transferred through capillary action (driven by the osmotic differential), before active processes are initiated in the stigma; for example, activation of PIPs by pH changes and phosphorylation to increase water transport (Liu et al. 2025; Mukherjee et al. 2024; Nieto-Giraldo et al. 2026).

Rozier et al. (2020) proposed that the SI response, in addition blocking pollen hydration, has a second checkpoint that inhibits pollen germination in the roughly 20% of SI pollen grains that hydrated similarly to compatible pollen. Perhaps it is the combined action of these two checkpoints that keeps SI pollen hydration at a very low rate and prevents pollen germination. One of the events that occurs rapidly after pollination is the regulation of stigma ROS levels in the stigma where stigma ROS levels are reduced following compatible pollinations and increased with SI pollinations (Huang et al. 2023; Liu et al. 2021). Focusing on the stigma PIPs as a key target for preventing pollen hydration, the increased stigma ROS with SI pollen could have inhibitory effects on the PIPs. The predominant ROS, hydrogen peroxide (H_2_O_2_), can inhibit PIP water transport so that H_2_O_2_ is transported by the PIPs into the cytoplasm. H_2_O_2_ was also found to cause PIPs to be internalized which would result in fewer PIPs at the plasma membrane for water transport (Mukherjee et al. 2024; Nieto-Giraldo et al. 2026). Previous studies have highlighted the importance of the vesicle trafficking through the exocyst and SNARE complexes in the stigma for promoting pollen hydration (Macgregor et al. 2023; Safavian et al. 2015; Samuel et al. 2009). Vesicle trafficking in the stigma is disrupted with SI pollen (Goring 2017; Macgregor et al. 2022; Samuel et al. 2009; Zhang et al. 2024) and perhaps this impacts PIP membrane dynamics to reduce water transport (Chevalier and Chaumont 2015). If there are more passive water transfer processes occurring in the early phase of compatible pollen hydration, then these SI signaling events would also be predicted to somehow disrupt these processes to account for the observed pollen hydration gap observed between the compatible pollen hydration mutants and SI pollen hydration.

## Suppl. Data

**Fig. S1.** Schematic of mutations in the tandemly-linked *MSL7* and *MSL8* genes.

**Table S1.** Candidate gene expression levels in the stigma.

**Table S2.** gRNA primers used to generate CRISPR/Cas9 vectors for MSL7 gene deletion mutations.

**Table S3.** Genotyping Primers.

## Acknowledgements

We thank work-study students (Liz Lo, Gary Chatha, Erin Navaratnam, Hamna Ammar and Cecilia Widjaja) for their technical assistance. We are also very grateful to Dr. James Doughty (University of Bath) for the *pcp-bα*/*β*/*γ* mutant seeds, Dr. Elizabeth Haswell (Washington University in St Louis) for the *msl7/8* mutant seeds, Dr. Anton Schäffner (Helmholtz Zentrum München) for the *pip* mutant seeds, and the ABRC for the *sd-rlk28* T-DNA mutant seeds (SALK_065560). This work was supported by a grant from the Natural Sciences and Engineering Research Council of Canada (NSERC) to DRG (RGPIN-2024-03945). PC was supported by an Aiken-Woods Memorial Scholarship.

## Author Contributions

All authors designed the research; PC and RL performed the research; PC and DG wrote the first draft; All authors analyzed the data, edited and approved the final version of the manuscript.

## Data availability

All data supporting the findings of this study are available within the paper and within its Suppl. data published online.

