## Supplemental Files for "Assessing the relative contributions of different *Arabidopsis thaliana* stigma factors to pollen hydration"

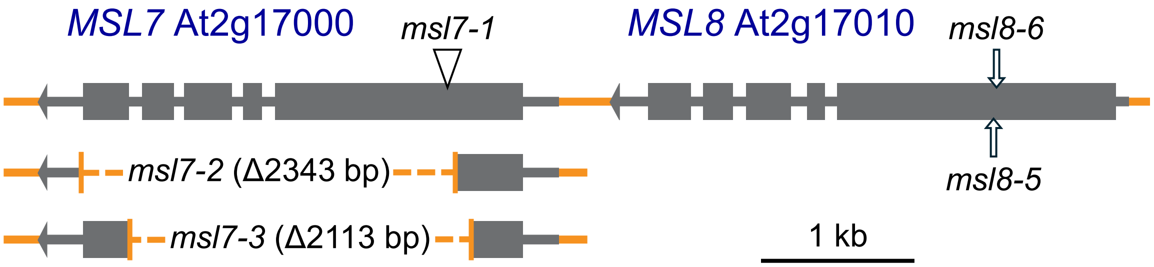

**Figure S1** Schematic of mutations in the tandemly-linked *MSL7* and *MSL8* genes. Dark grey boxes represent exons, and light grey line represents introns and non-coding sequences. The triangle represents the location of the T-DNA insertion in *msl7-1* (SALK_133223; Wang et al., 2022). The orange dotted lines represent the *MSL7* gene deletion mutations for *msl7-2* and *msl7-3* (this paper). The arrows on the first exon of *MSL8* represent the location of the *msl8-5* and *msl8-6* mutations (*msl8-5* 2 bp insertion + 13 bp deletion, *msl8-6* 1 bp insertion; Wang et al., 2022).

**Table S1. Candidate gene expression levels in the stigma.**

| | | Stigmatic Papillae (T1 Average) | Stigmas (TRAVA) | |
|---|---|---|---|---|
| *AtPIP1;1* | At3g61430 | 37 | 487 | * |
| *AtPIP1;2* | At2g45960 | 1365 | 23599 | * |
| *AtPIP1;3* | At1g01620 | 16 | 1600 | |
| *AtPIP1;4* | At4g00430 | 352 | 3916 | |
| *AtPIP1;5* | At4g23400 | 13 | 772 | |
| *AtPIP2;1* | At3g53420 | 59 | 1222 | * |
| *AtPIP2;2* | At2g37170 | 2237 | 5384 | * |
| *AtPIP2;3* | At2g37180 | 696 | 1569 | |
| *AtPIP2;4* | At5g60660 | 0 | 186 | |
| *AtPIP2;5* | At3g54820 | 544 | 5582 | |
| *AtPIP2;6* | At2g39010 | 254 | 1054 | |
| *AtPIP2;7* | At4g35100 | 887 | 6985 | |
| *AtPIP2;8* | At2g16850 | 0 | 77 | |
| *MSL7* | At2g17000 | 1035 | 9530 | |
| *MSL8* | At2g17010 | 8 | 7 | |
| *RKF1* | At1g29750 | 39 | 1056 | |
| *RKFL3* | At1g29740 | 0 | 25 | |
| *RKFL2* | At1g29730 | 4 | 62 | |
| *RKFL1* | At1g29720 | 6 | 697 | |
| *SD-RLK28* | At4g00340 | 228 | 2529 | |
| *PCP-Bα* | At5g61605 | 0 | 0 | |
| *PCP-Bβ* | At2g29790 | 0 | 1 | |
| *PCP-Bγ* | At2g16535 | 0 | 0 | |
| *FERONIA* | At3g51550 | 667 | 7036 | |
| *ANJ* | At5g59700 | 84 | 1380 | |
| *CVY1* | At2g39360 | 154 | 275 | |
| *HERK1* | At3g46290 | 51 | 694 | |

Other known pollen regulators in the stigmatic papilla added as a comparison (CrRLK1L-1 receptor kinases)

| Stigmatic Papillae (T1 Average) | Stigmas (TRAVA) |
|---|---|
| Scale | Scale |
| 0 | 0 |
| 200 | 1000 |
| 400 | 2000 |
| 600 | 3000 |
| 800 | 4000 |
| 1000 | 5000 |
| 1200 | 6000 |
| 1400 | 7000 |
| 1600 | 8000 |
| 1800 | 9000 |
| >2000 | >10000 |

*Quad pip mutant

*pip1;1-1*
*pip1;2-2*
*pip2;1-2*
*pip2;2-3*

**Table S2. gRNA primers used to generate CRISPR/Cas9 vectors for MSL7 gene deletion mutations.**

| Gene | | Sequence |
|---|---|---|
| MSL7 At2g17000 (this study) | msl7-2 gRNA Forward Primer | ATATATGGTCTCGATTGGATAGATCCGCCACAAGAAGGTTTTAGAGCTAGAAATAGCAAGTTAAAAT |
| | msl7-2 gRNA Reverse Primer | ATTATTGGTCTCTAAACTCAGGGTTTTGTGACCAGGCCAATCTCTTAGTCGACTCTACCA |
| | msl7-3 gRNA Forward Primer | ATATATGGTCTCGATTGAGAGCTAGTTTCGAGGGCGAGTTTTAGAGCTAGAAATAGCAAGTTAAAAT |
| | msl7-3 gRNA Reverse Primer | ATTATTGGTCTCTAAACATACTCTGGCTTGCTATCAACAATCTCTTAGTCGACTCTACCAATA |

**Table S3. Genotyping Primers.**

| Genotyping Function | Primer Name | Primer Sequence (5' to 3') | Reference |
|---|---|---|---|
| *msl7-2, msl7-3* CRISPR mutants | msl7 del LP | ACTTTGTTCCGCTGCGTTTC | this study |
| | msl7 del RP | TGAACAAGAAGTAGTACCAGCCA | |
| wildtype *MSL7* allele (At2g17000) | msl7 int LP | AAGCACCGGAGATCAAAACG | this study |
| | msl7 int RP | TCCCACTCTTTGCTTTCACC | |
| msl8-5 (At2g17010) | CRISPR/Cas9 mutation in exon 1 | | *msl8-5* single mutant |
| *msl8-6* (At2g17010) | CRISPR/Cas9 mutation in exon 1 | | *msl7-1 msl8-6* double mutant |
| *msl7-1* T-DNA mutant (SALK_133223) | LBb1.3 | ATTTTGCCGATTTCGGAAC | |
| | SALK_MSL7 RP | GAATTCCGCAAACCCTTTAAG | Wang et al. (2022) |
| *MSL7* wildtype allele (At2g17000) | SALK_MSL7 LP | AAATCAGCTCCTCCTTTCTGC | doi.org/10.1093/jxb/erab525 |
| | SALK_MSL7 RP | GAATTCCGCAAACCCTTTAAG | |
| *rkfΔ-1, rkfΔ-2* CRISPR deletion mutants | RKF1 cluster del LP | TTGATGATTGTTAAGGCCAAGAAG | *RKF1* gene cluster deletions |
| (At1g2970, At1g29730, At1g29740, At1g29750) | RKF1 cluster del RP | GGTTACCCAATTGATAAGCGAG | (*RKF1 RKFL1-3* ) |
| *RKF1* wildtype allele (At1g29750) | RKF1 internal LP | TCGCTGAGATACGGTTCACC | Lee and Goring (2021) |
| | RKF1 internal RP | GCACGGCTTGGAATCTGTAC | doi.org/10.1093/jxb/eraa496 |
| *sd-rlk28* T-DNA mutant (SALK_065560 ) | LBa1 | TGGTTCACGTAGTGGGCCATCG | Guo et al. (2023) |
| | SD-RLK28 RP | CTTAGAAAGACCTGGAAGCGG | doi.org/10.1111/jipb.13547 |
| *SD-RLK28* wildtype allele (At4g00340) | SD-RLK28 LP | TGGGTTTATTGATTCAATCACG | |
| | SD-RLK28 RP | CTTAGAAAGACCTGGAAGCGG | |
| *pcp-bα-1* T-DNA mutant (SALK_207087) | LBb1.3 | ATTTTGCCGATTTCGGAAC | *pcp-bα/β/γ* triple mutant |
| | PCP-Bα RP | TTTATCAAGAAAACAGGTCCTCGG | |
| *PCP-Bα* wildtype allele (At5g61605) | PCP-Bα LP | TGCCTCTTCATGATTTTCCTCGTTCC | Wang et al. (2017) |
| | PCP-Bα RP | TTTATCAAGAAAACAGGTCCTCGG | doi.org/10.1111/nph.14162 |
| *pcp-bβ-1* T-DNA mutant (SALK_062825) | LBb1.3 | ATTTTGCCGATTTCGGAAC | |
| | PCP-Bβ RP | CGGTTGCAGACAGCCACACAC | |
| *PCP-Bβ* wildtype allele (At2g29790) | PCP-Bβ LP | GTAGTTTCTCTCGTTCCTCATGG | |
| | PCP-Bβ RP | CGGTTGCAGACAGCCACACAC | |
| *pcp-bγ-1* T-DNA mutant (SALK_072366) | LBb1.3 | ATTTTGCCGATTTCGGAAC | |
| | PCP-Bγ RP | TTGCAGTCAGCAACACATTCTG | |
| *PCP-Bγ* wildtype allele (At2g16535) | PCP-Bγ LP | CATCATCCTCATTTCTTCATTTCC | |
| | PCP-Bγ RP | TTGCAGTCAGCAACACATTCTG | |
| *pip1;1-1* T-DNA mutant (GABI_437B11) | GABI_LB2 | CCCATTTGGACGTGAATGTAGACAC | *pip1;2-2, pip2;1-2, pip2;2-3* single mutants |
| | pip1;1-1 RP | ACAGACCAAAAGTAACCGC | |
| *PIP1;1* wildtype allele (At3g61430) | pip1;1-1 LP | TTATCACTGACCCAATCCA | *pip1;1 pip1;2 pip2;1 pip2;2* quadruple mutant |
| | pip1;1-1 RP | ACAGACCAAAAGTAACCGC | |
| *pip1;2-2* T-DNA mutant (SALK_019794) | LBb1.3 | ATTTTGCCGATTTCGGAAC | |
| | pip1;2-2 RP | AGTTGCCTGCTTGAGATAAACC | Ceciliato et al. (2019) |
| *PIP1;2* wildtype allele (At2g45960) | pip1;2-2 LP | AGTTCACTGGTTTCTCCGAT | 10.1186/s13007-019-0423-y |
| | pip1;2-2 RP | AGTTGCCTGCTTGAGATAAACC | Schley et al. (2025) |
| *pip2;1-2* T-DNA mutant (SM_3_35928) (CS122639) | Spm32 | TACGAATAAGAGCGTCCATTTTAGAGTGA | doi.org/10.1111/pce.15222 |
| | pip2;1-2 RP | CAACGCATAAGAACCTCTTTGA | |
| *PIP2;1* wildtype allele (At3g53420) | pip2;1-2 LP | CTAACCACTCCAACAGAGAAG | |
| | pip2;1-2 RP | CAACGCATAAGAACCTCTTTGA | |
| *pip2;2-3* T-DNA mutant (SAIL_169_A03) (CS871747) | SAIL LB3 | TAGCATCTGAATTTCATAACCAATCTCGATACAC | |
| | pip2;2-3 RP | TTATAGATTACGGCAGCTCCG | |
| *PIP2;2* wildtype allele (At2g37170) | pip2;2-3 LP | CAACCATAAGCCTACCAAAAGG | |
| | pip2;2-3 RP | TTATAGATTACGGCAGCTCCG | |
